# Human Proteome Project Mass Spectrometry Data Interpretation Guidelines 3.0

**DOI:** 10.1101/733576

**Authors:** Eric W. Deutsch, Lydie Lane, Christopher M. Overall, Nuno Bandeira, Mark S. Baker, Charles Pineau, Robert L. Moritz, Fernando Corrales, Sandra Orchard, Jennifer E. Van Eyk, Young-Ki Paik, Susan T. Weintraub, Yves Vandenbrouck, Gilbert S. Omenn

## Abstract

The Human Proteome Organization’s (HUPO) Human Proteome Project (HPP) developed Mass Spectrometry (MS) Data Interpretation Guidelines that have been applied since 2016. These guidelines have helped ensure that the emerging draft of the complete human proteome is highly accurate and with low numbers of false-positive protein identifications. Here, we describe an update to these guidelines based on consensus-reaching discussions with the wider HPP community over the past year. The revised 3.0 guidelines address several major and minor identified gaps. We have added guidelines for emerging data independent acquisition (DIA) MS workflows and for use of the new Universal Spectrum Identifier (USI) system being developed by the HUPO Proteomics Standards Initiative (PSI). In addition, we discuss updates to the standard HPP pipeline for collecting MS evidence for all proteins in the HPP, including refinements to minimum evidence. We present a new plan for incorporating MassIVE-KB into the HPP pipeline for the next (HPP 2020) cycle in order to obtain more comprehensive coverage of public MS data sets. The main checklist has been reorganized under headings and subitems and related guidelines have been grouped. In sum, Version 2.1 of the HPP MS Data Interpretation Guidelines has served well and this timely update to version 3.0 will aid the HPP as it approaches its goal of collecting and curating MS evidence of translation and expression for all predicted ∼20,000 human proteins encoded by the human genome.

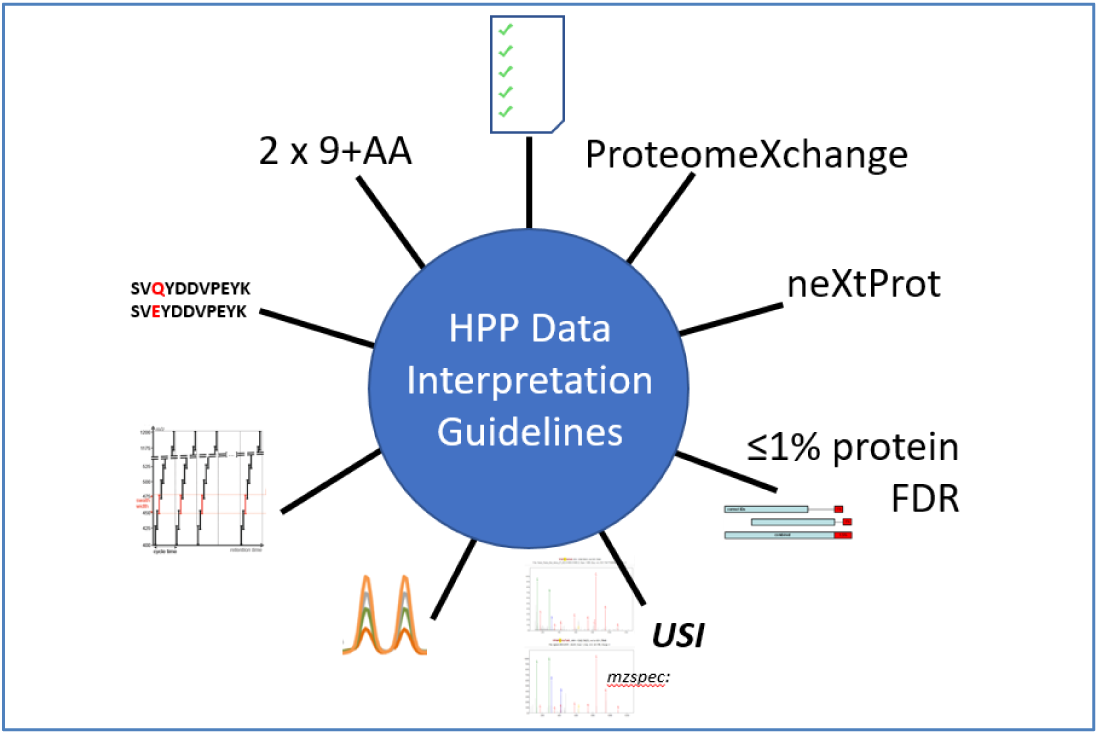

## Introduction

The Human Proteome Organization’s^1^ (HUPO) Human Proteome Project^2,3^ (HPP) was launched in 2010 as an international endeavor to build on the success of the Human Genome Project^4,5^ by characterizing the products of the ∼20,000 human protein-coding genes. As of January 2019, 17,694 proteins demonstrated compelling mass spectrometry (MS) or non-MS protein-level evidence in neXtProt (i.e., PE1), leaving 2129 proteins without strong evidence (PE2,3,4) that were have been designated as the HPP’s ‘missing proteins’^6^. The PE2,3,4 missing proteins represented 10.7% of all neXtProt’s PE2,3,4 proteins. The goals of the HPP are (1) to complete the protein ‘parts’ list, including isoforms, post-translational modifications (PTMs), and single amino acid variants, with characterization of their functions; and (2) to make proteomics an integral part of all multi-omics studies in life sciences. The Chromosome-centric HPP (C-HPP) consortium focused largely, but not exclusively, on the first two goals^7^, whereas the Biology and Disease HPP (B/D-HPP) focused largely on the latter goal, whilst recognizing that many studies will also uncover disease-specific or tissue-specific PE2,3,4 missing proteins. The progress in achieving these goals has been tracked yearly via a set of published metrics^3,8–11^ based on the major knowledge bases of the HPP, namely neXtProt^12^, PeptideAtlas^13–15^, Human Protein Atlas^16^, and the ProteomeXchange^17,18^ consortium of proteomics data repositories.

In order to maintain a high standard of quality for the identifications in the compendium of human proteins, and to ensure that journal articles and data contributions are reporting with equally high standards, a set of MS data guidelines was developed. The inaugural version 1.0 guidelines were released in 2013 and mandated deposition of data to members of the newly formed ProteomeXchange Consortium for proteomics/MS data and other repositories for other kinds of biochemical data. Initial progress in protein detection was rapid since there were many high abundance proteins present in common samples available to catalog. However, it soon became apparent that, as the compendium of proteins commonly seen in high abundance became complete, the control of false positives during the hunt for missing proteins became imperative. Version 2.1 of the HPP MS Data Interpretation Guidelines was developed and published in 2016^19^. These guidelines went beyond data deposition requirements, setting out minimum standards for the handling of false discovery rate (FDR) in the interpretation of MS data as well as minimum standards to claim the detection of any missing protein or protein otherwise not yet found in the HPP KB compendium of detected proteins.

Version 2.1 of the guidelines has been in force for the annual Journal of Proteome Research HPP Special Issues beginning in 2016. For papers submitted outside the frame of the Special Issue, the Editors of the Journal of Proteome Research and increasing numbers of other proteomics journals now require these guidelines to be met for all claims of missing protein identifications.

In the past year it has become apparent that, despite the advances in proteomics, the increased difficulty of detecting the remaining ∼10% of the human proteome requires an update to the guidelines, as discussed on-line (https://docs.google.com/document/d/167wLMYshQ3jUPJonxyk6TcOvT8GrZcqYOZAtUGOK29w), in the Bioinformatics Hub at the 2018 17^th^ HUPO World Congress in Orlando, USA, at the 2019 21^st^ C-HPP Symposium in Saint-Malo, France, and at the 2019 18^th^ HUPO World Congress in Adelaide, Australia. In these venues, the leadership of the HPP, along with other interested contributors, debated 25 aspects of the existing guidelines for journal articles as well as current practices of the pipelines that maintain and refine the resources that comprise the HPP KB ecosystem.

Here, we describe the outcomes of these discussions, which are reflected in a refined set of guidelines to take the HPP forward. First, we present a revised version 3.0 of the HPP MS Data Interpretation Guidelines in the form of a brief one-page checklist and more extensive three-page checklist documentation. Next, we discuss the reasoning behind the changes to each guideline, often providing the set of options debated. Finally, we discuss the reasoning behind the changes to overall HPP policy used by the HPP KB pipeline that tracks and disseminates the best-available gathered understanding of the human proteome as the HPP and the global community gear up to tackle the most difficult refractory proteins of the human proteome.

## Changes to the guidelines

Whilst version 3.0 of the checklist (Supplementary Material S1) looks similar to the previous version 2.1 (https://www.hupo.org/HPP-Data-Interpretation-Guidelines), it contains major differences. First, in addition to the requirement of a checkmark to indicate that each requirement is fulfilled (or NA for not applicable or NC for not completed, both of which require an explanation as to why this is not applicable or complete at the bottom of the checklist), a new column requests a location where the pertinent information may be found. This will typically be a reference to a page number or a supplementary document. This additional information makes it much easier for the journal editors, reviewers, and readers to find the section in the submission that fulfills each guideline, which can sometimes be difficult and slows reviewing. Such a requirement for page numbers is already common in submission checklists for many bioinformatics journals.

A second substantial difference is the reorganization of the guidelines into numbered major items and lettered subitems. Whereas the previous version had 15 major items, some of which were highly related and needed to be considered together, the latest version has only 9 major items, but some of those contain subitems that should be considered as a group. We hope that this provides a better overall organizational framework and is more user-friendly ensuring contributor completion. A few guidelines have been deemphasized by being merged with important related guidelines. Two new guidelines have been added, as discussed in detail below. In the following paragraphs each guideline will be discussed briefly with emphasis on changes since version 2.1.0. There are two parts of the guidelines: reports of well-established proteins and reports of claims of novel detection of predicted proteins.

### Section 1

Guideline 1 remains essentially unchanged as a formal requirement that each manuscript be submitted with a filled-in checklist describing the compliance of the manuscript with the guidelines. If any manuscript is received for publication without a checklist, the handling editors immediately request this be completed before sending any manuscript for review. The extended description of Guideline 1 has been augmented to describe the new requirement in column two for a page number with line number or paragraph number, or other indication of the location (such as a specific supplemental table) of the requested information.

Guideline 2 requiring deposition of all data sets into a ProteomeXchange repository has been expanded into four subparts because, in the version 2.1 guidelines, these subparts were concatenated into one sentence, where compliance suffered. There was a strong tendency for authors to fulfill the first part of the requirement and move on without addressing other components. In order to avoid this, the four main aspects of the previous guideline 2 have been separated into four subitems 2a, b, c, d, which require complete data deposition, deposition of analysis reference files, PXD identifier in abstract, and reviewer credentials, respectively.

Guideline 3, requiring the use of the most recent neXtProt release, rather than older versions, remains unchanged. Our understanding of the human proteome continues to evolve rapidly and the use of older versions may lead to confusion and outdated claims. Generally, neXtProt curators update regularly, with their prior January release relied upon for HPP Journal of Proteome Research Special Issue data analysis/reanalysis and which effectively is reflected as the annual HPP metrics^3,6,9–11,20^.

Guideline 4 merges all previous FDR-related guidelines (4 – 9 in version 2.1) into a single top-level entry with four subitems designed to streamline this section. Previous top-level guidelines 7 and 8 request that authors consider that FDR calculations should be reported with an appropriate number of significant digits (usually one or two), because they are based on several imperfect assumptions, and that the required FDR calculations and implied number of wrong identifications should be carefully considered in later analysis of any resulting protein list. These points have been merged into part b of guideline 4, which also addresses reporting of FDR values. The HPP community seems to understand these aspects well and separate items no longer seem necessary.

### Section 2

Whereas guidelines 1 to 4 apply to all manuscripts presenting MS data, the following guidelines 5 to 9 apply only to manuscripts presenting evidence that could qualify the newly-identified proteins for consideration as PE1 in neXtProt or to provide MS evidence for PE1 proteins lacking MS data, so classified based on other types of data.

In the previous version of the guidelines, missing protein MS evidence was referred to as “extraordinary detection claims”, reminiscent of the aphorism that “extraordinary claims require extraordinary evidence”, often credited to Carl Sagan (https://en.wikipedia.org/wiki/Sagan_standard) or Amos Bairoch (https://en.wikipedia.org/wiki/Amos_Bairoch). The phrase “extraordinary detection claims” was confusing to many, so this phrase has been replaced by “claims of new PE1 protein detection”. Such claims may apply to one of the “missing proteins” currently in neXtProt with protein existence status of PE2,3,4. They may apply to a current PE5 protein, although most of these entries are annotated as pseudogenes in UniProtKB and additional care should be applied to justify that their detection is not merely a variation of the common PE1 protein that the predicted PE5 protein sequences closely resemble. Finally, this assignation may apply to a protein not yet listed in neXtProt. These might include: (i) an entry not yet manually reviewed in UniProtKB, (ii) a protein currently annotated as a lncRNA, (iii) a smORF, or (iv) some other novel protein-coding element. There are many new protein entries, including immunoglobulins, in annual releases from neXtProt. The first three guidelines are specific to each of three different acquisition technologies, whereas the two guidelines that follow apply to all three technologies—DDA, SRM, and DIA-MS.

Guideline 5 has become a guideline containing three subitems that merge several previous top-level guidelines into a single one focused on requirements for data dependent acquisition (DDA) MS workflows, commonly referred to as “shotgun proteomics”. Part 5a is essentially the same as previous guideline 10, which affirms that evidence spectra for new PE1 protein detection claims must be high mass-accuracy, high signal-to-noise ratio, and clearly annotated with peak interpretations. The previous guideline 11 enjoining authors to examine the spectra carefully for telltale signs of misidentification has been appended to the extended description of the new subitem “a”, since, although laudable, it was difficult for many authors to perform effectively and is less important in the presence of the guideline requiring comparison with a synthetic spectrum.

Guideline 5b is similar to the previous guideline 12, seeking clear presentations of synthetic peptide spectra that match endogenous peptide spectra. The guideline has been augmented to include a recommendation by the guidelines revision team group that spectra derived from digested recombinant proteins are suitable substitutes for those MS spectra derived from peptides created with peptide synthesizing technologies. The guideline has also been amended so that a retention time match between the target and the synthetic peptide are no longer required, but rather suggested only if the target and reference are both run on the same instrument. The use of public reference spectra from synthetic peptides such as from SRMAtlas^21^ is now specifically allowed.

Guideline 5c is <u>completely</u> new. A persistent problem with discussions about the merits of certain spectra as evidence for new PE1 protein detection claims is the general inability to identify specific spectra and access them easily in the data repositories for close examination. PDF representations of MS spectra found in supplementary materials are useful but resist close examination and the application of KB tools that reviewers or readers might like to use for inspection of presented MS evidence. Furthermore, if reprocessing of the data set does not yield the same result, it is very difficult to assess what became of the key spectrum and why it does not reveal the same PSM in reprocessing. In order to solve this problem, the HUPO Proteomics Standards Initiative^22–24^ (PSI) has developed the Universal Spectrum Identifier (USI) concept as a multi-part key that can universally identify any acquired spectrum in a manner that any repository containing the data set would be able to display or furnish the same spectrum via this identifier. Guideline 5c now introduces a requirement for the provision of USIs for all spectra that provide evidence for new PE1 protein detection claims, natural sample observations and synthetic peptide spectra alike. See http://psidev.info/USI for more information on how to create and use USIs.

Guideline 6 is the same as guideline 13 in version 2.1. It applies to selected/multiple reaction monitoring (SRM/MRM) workflows^25^, requiring that chromatogram traces of the detected peptides be provided along with the matching chromatograms of heavy-labeled reference synthetic peptides. It is important in SRM that both the intensity patterns and the retention times match, since there are typically far fewer ions monitored than peaks available in full spectra. We have added a request that the heavy-labeled reference peptides should be spiked in at an abundance similar to the target peptides so that minor impurities in the reference do not contribute to the target signal. If the heavy-labeled spike-in has a 1% impurity in the form of light peptide, then, if the reference is spiked in at 100 times the target peptide abundance, the impurity will contribute as much signal as the target peptide, leading to an incorrect abundance or even a spurious detection. The extended description reaffirms that guidelines 8 and 9 also apply to SRM as there has been some confusion previously. This same guideline can also be applied to parallel reaction monitoring^26^ (PRM) data, although since PRM acquisition creates full MS/MS spectra, full compliance with guideline 5 is also acceptable.

Guideline 7 is a new guideline that addresses untargeted data independent acquisition (DIA) workflows such as SWATH-MS^27^ and similar techniques^28^. The version 2.1 guidelines were silent on DIA data sets as we felt that the technology was too new to write useful guidelines at that time. In the meantime, DIA has become a broadly-adopted technology. Although DIA has not yet been used to claim detection of new PE1 proteins, this will surely come. Guideline 7 is simple. It applies guidelines 5 and 6 depending on the mode of bioinformatics analysis of the DIA data. If the data are analyzed via extracted ion chromatograms (XICs) (sometimes called peptide-centric analysis) such as with OpenSWATH^29^, Spectronaut^30^, and SWATH 2.0, then the SRM guideline 6 applies. If the data are analyzed via extracted demultiplexed spectra (sometimes called spectrum-centric analysis) such as with DIA-Umpire^31^ and DISCO (Shteynberg et al, in preparation), then the DDA guidelines 5a-c apply. The next few years will show whether DIA can be used reliably for new PE1 detection claims and if this simple approach to a DIA guideline is sufficient. Of interest is the observation that the journal *Molecular and Cellular Proteomics* has recently developed a comprehensive set of guidelines for handling DIA data^32^. Authors are advised to examine these and use these where applicable to further support claims, although as yet they are not required as part of the HPP and/or *Journal of Proteome Research* guidelines.

Guideline 8 remains the same as the previous guideline 14, encouraging authors to consider alternate explanations for novel spectral matches. In many cases a single amino acid variant (SAAV) or a post-translational modification (PTM) creates an isobaric or near-isobaric change that can mean the difference between mapping to a protein never before detected with MS and a common protein observed by millions of PSMs. Despite some useful tools available for authors to address this guideline (e.g., neXtProt peptide uniqueness checker^33^ and PeptideAtlas ProteoMapper^34^), it remains one of the most difficult to fulfill, since exact mappings are clear enough, but near mappings are difficult and time consuming to assess. Nevertheless, the authors consider that this remains an important guideline that researchers and reviewers should continue to consider when presenting new PE1 protein detection claims.

Guideline 9 is a derivative of the previous guideline 15, although many aspects were discussed extensively and several small modifications made. This guideline provides the minimum MS requirements for the number and attributes of peptides that support the claim of any new PE1 protein detection. The group reaffirmed that two uniquely-mapping, non-nested peptides of nine or more amino acids should be the minimum required for such a claim. However, various aspects of this requirement were discussed extensively and clarifications made. First, the definition of non-nesting was clarified. Strictly, the meaning of non-nested means that one peptide may not be completely contained within another. The reasoning is that, although the observation of two nested peptides increases the confidence that the peptides have been correctly identified, it does not provide any additional evidence that the peptide has been correctly mapped to the protein in question; i.e., if the longer peptide is mismapped and should instead map to some part of the proteome that we do not yet fully understand (such an immunoglobulin or some other variation), then the nested peptide will have exactly the same problem, and provides no new information. An extension of one peptide beyond the other provides some additional mapping confidence. However, the previous guidelines as written permitted even a single amino acid extension. For example, a tryptic peptide PEPTIDESR and a LysargiNase^35^ (that cleaves before K/R instead of after K/R) peptide KPEPTIDES would qualify as non-nested under the previous guidelines. We recommend amendment of the guidelines to require that the total extent of the coverage of the two nested peptides combined be at minimum 18 amino acids (2 × 9). This strategy had already been implemented at neXtProt, and thus there is no change there, but does reflect a change for PeptideAtlas and other interpretations of the guidelines.

Guideline 9 now also contains a clarification for how to handle identical proteins. There are 118 entries in the current January 2019 neXtProt release that have the same protein sequence for at least one other entry. These entries reflect different gene loci that may have synonymous-coding nucleotide variation but yield the exact same protein sequence. This is an extreme case of highly homologous proteins. These 118 entries can be retrieved by applying the SPARQL query NXQ_00231 (https://www.nextprot.org/proteins/search?mode=advanced&queryId=NXQ_00231) in the neXtProt advanced search tool. They represent a total of 51 distinct protein sequences. It was decided that for the purposes of PE status assignment, if two or more qualifying peptides map uniquely to multiple identical proteins, then all such proteins will be switched to PE1 as a group since they are indistinguishable from each other. Nonetheless, it was noted that since their gene promoter regions are likely to differ, these proteins may be expressed in different tissues, or under different spatiotemporal circumstances or under different physiological or pathological conditions. As is the case now, each will be counted individually as PE1.

The group further clarified that, while the two peptides presented as evidence do not need to originate from the same sample or instrument, they do need to be presented together in the paper. The practice of offering a single new suitable peptide to complement a pre-existing different suitable peptide already in PeptideAtlas and neXtProt is permitted, but the PeptideAtlas peptide spectrum must also be scrutinized and compared with a synthetic peptide spectrum in accordance with the above guidelines with all evidence presented in the paper.

### Changes to the HPP PE2,3,4 missing protein strategy

The current basic process by which the HPP investigators manage the process of reducing the number of missing proteins of the human proteome, herein called the “HPP pipeline”, begins with the collection of MS data sets from the global community and deposition in one of the ProteomeXchange repositories. The vast majority of data sets are deposited into PRIDE^36,37^, with some routed through MassIVE^38^ and jPOST^39^. These data sets may come from experiments presented in HPP special issues such as this issue, or from experiments performed by other members of the community in pursuit of their own research objectives. After ProteomeXchange deposition, PeptideAtlas collects raw MS data files and reprocesses those data using the tools of the Trans-Proteomic Pipeline (TPP)^40–42^. Thresholds are set extremely high in PeptideAtlas in order to obtain a 1% protein-level FDR across the ensemble of all data sets. In November each year, PeptideAtlas stops processing new data sets and creates an annual build reflecting the current state of the human proteome from MS evidence. In December the final peptide list is transferred to neXtProt for integration into neXtProt’s next build/release based on their import of the most recent version of the human proteome from UniProtKB/Swiss-Prot. While all peptides that pass thresholds are visible in PeptideAtlas and neXtProt, only the proteins with two uniquely-mapping non-nested peptides with length 9 amino acid (AA) or greater, as called by neXtProt, are deemed to have sufficient evidence to be labeled as confidently detected PE1 proteins by MS methods. Therefore, neXtProt is the final arbiter to decide if a PE2,3,4 protein in UniProtKB is deemed PE1 in neXtProt and released as such for HUPO’s HPP. Figure 1 provides a graphical summary of the current HPP pipeline.

**Figure 1.**
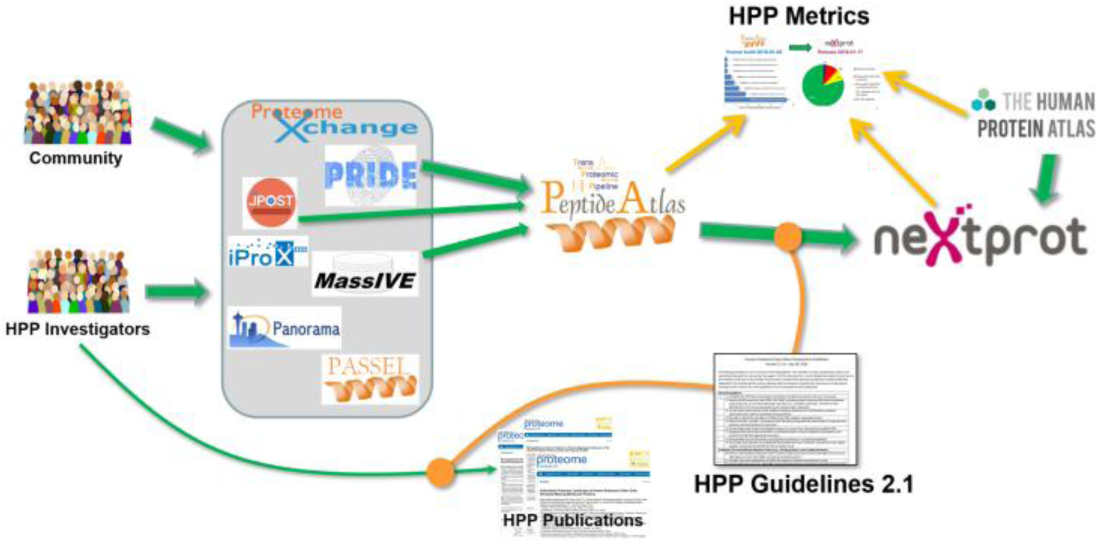
Overview of the 2019 HPP pipeline for data integration. HPP investigators publish their results constrained by the HPP guidelines. The data sets from these publications as well as other data sets from the community flow into the ProteomeXchange repositories. Currently a subset of the data sets from PRIDE, MassIVE, and JPOST are reprocessed by PeptideAtlas, the results of which are transferred to neXtProt constrained by the HPP guidelines. Information from PeptideAtlas, neXtProt, and Human Protein Atlas is summarized yearly in the HPP Metrics summary (this issue). Data from the Human Protein Atlas is also transferred to and reprocessed by neXtProt as part of the HPP data cycle, although they are not yet used to change PE status.

The group discussed several ambiguities and refinements of this process and made recommendations/decisions on how the HPP pipeline will be defined for the next few years. The group also sought to clarify some terminology pertaining to peptides relevant to the HPP Pipeline process, most notably the term “stranded peptide”, which has been used in several different (possibly confusing) contexts in the past^43,44^. After considerable discussion, it was resolved that the term “stranded peptide” shall specifically refer to “a peptide that meets the minimum length and mapping uniqueness requirements and has publicly available evidence for its detection via MS, but the evidence is not within the HPP Pipeline”. In order to become unstranded, this publicly available information must be captured and validated by the HPP Pipeline. In addition, the term “singleton peptide” shall refer to a peptide that meets the minimum length and mapping uniqueness requirements but does not have the needed additional partner peptide to achieve the full requirements for two non-nested peptides. Stranded peptides may be singletons or not; singleton peptides may be stranded or not. This terminology is used further below.

The first refinement is for how SAAVs are handled with respect to mapping uniqueness. The fundamental question is what degree of mutation should be considered when mapping potentially uniquely-mapping peptides to the proteome. Should all SAAVs in neXtProt be considered when mapping peptides, and in all permutations (e.g., if there are three annotated mutation sites in a single peptide, should mapping all three residues that are mutated be considered, or should just one at a time be considered)? Despite substantial diversity in opinion, the consensus was that co-occurrence of nearby SAAVs was very low, and therefore simply considering one mutation per peptide was sufficient. All mutations in neXtProt, except for the somatic mutations from COSMIC^45^, will be considered during mapping of peptides to proteins.

The group discussed whether there should be some formal adjustment to the lower limit requirement of two peptides of nine amino acids or longer. These requirements are designed to ensure a certain level of confidence in the MS detection of missing proteins, but this level was never really quantified in a way to justify that 2 × 9 should be sufficient, but (2 × 8) or (3 × 8) or (1 × 9) + (1 × 8) + (1 × 7) should not. An example of the latter comes in the form of the current state of the protein Q8N688 □-defensin 123, which has multiple detections of 7 AA, 8 AA, and 9 AA peptides as shown in Figure 2 and is claimed by Wang et al^46^. Because this protein is only 67 amino acids long, and the mature form is only 47 AAs long after cleaving off the 20 AA-long signal peptide, these are the only three tryptic peptides that can be expected. The obvious question is: should this complete MS evidence be sufficient for PE1 status assignment? After substantial discussion, it was decided that there would be no change to the 2 × 9 policy for now, because building in a more intricate limit without a mathematical/statistical foundation for doing so was inadvisable. It was deemed that the 2 × 9 policy was simple and clear and worth retaining in the absence of a more compelling lower limit. However, two future courses of action were recommended.

**Figure 2.**
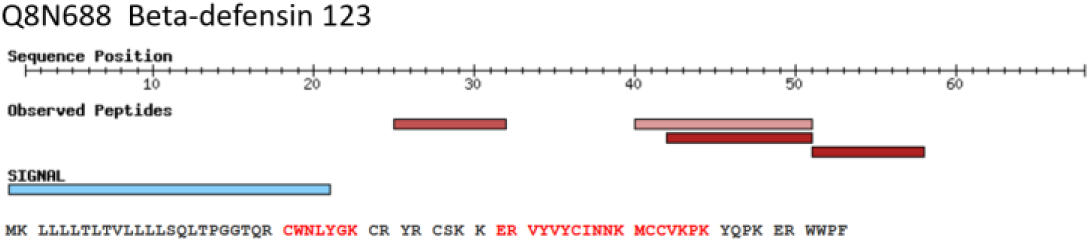
Depiction of the current status of Q8N688 Beta-defensin 123. The protein is only 47 AAs long after cleavage of the 20 AA signal peptide. Three distinct peptide sequences are detected (plus a fully nested peptide), but only one of the three meets guideline length requirements. Yet, all of the expected tryptic peptides (plus one missed cleavage product) are detected with excellent spectra. Should this be sufficient?

First, a sounder justification for the lower limit should be sought, perhaps one where a single probability formed the lower bound, and there might be multiple combinations that can achieve this probability. This would likely yield a per-protein metric since it is far easier and far more likely to obtain peptides that map to a very long protein than a very short one. In many cases, the use of multiple proteases might be needed to overcome the limitation of reliance on tryptic peptides, as might use of semi-tryptic and N or C-terminal peptides (see below and the 2018 HPP metrics^3^).

Second, guidelines v2.1 contained an “exceptions clause” for consideration of special cases. However, no mechanism was defined or implemented to deal with these special cases until now. The group recommended that a dedicated expert panel be formed by the HPP Knowledge Base Pillar Committee to judge whether particular proteins (including short proteins) that do not meet the guidelines precisely as written may indeed have sufficient evidence to meet the HPP’s desired level of confidence for PE1 status assignment. For each of the proteins recommended as candidates for elevation to PE1 without the minimum MS evidence, the panel would review the available spectra and prospects for obtaining additional MS evidence. In some cases, useful confirmatory non-MS evidence may exist. If the obtainable evidence is excellent despite not meeting the guidelines and further MS evidence is deemed unlikely, such proteins could be proposed by the panel to neXtProt for assessment as PE1. □-defensin 123 (gene name DEFB123) shown above was a prime initial exemplar candidate for the expert panel to consider.

The group discussed whether there are any proteins that should be declared too difficult and unachievable, and should therefore be simply removed from the denominator of the ratio describing the fraction of detectable proteins in the human proteome identified as PE1 proteins. As an example, there are 15 proteins which cannot generate two uniquely mapping 2 × 9 peptides even when using a series of five different common proteolytic enzymes (trypsin, LysargiNase, GluC, AspN and chymotrypsin). See Supplemental Table 1 for a list of these proteins. Should such proteins be declared unattainable with MS technologies? Remarkably, of these 15, nine are already designated as PE1, one of which (C9JFL3) has remarkably good non-protease-specific peptides from N and C termini as depicted in Figure 3. This is common when a protein is highly abundant in a sample. Thus, the group decided that no proteins would be declared too difficult now since, if enriched or purified to sufficient abundance, many might be accessed from the termini with the aid of non-specific cleavages (e.g., through carboxypeptidases). Enrichment of PTM-containing proteins, such as shown with SUMOylation^3,47^ may also be especially effective here.

**Figure 3.**
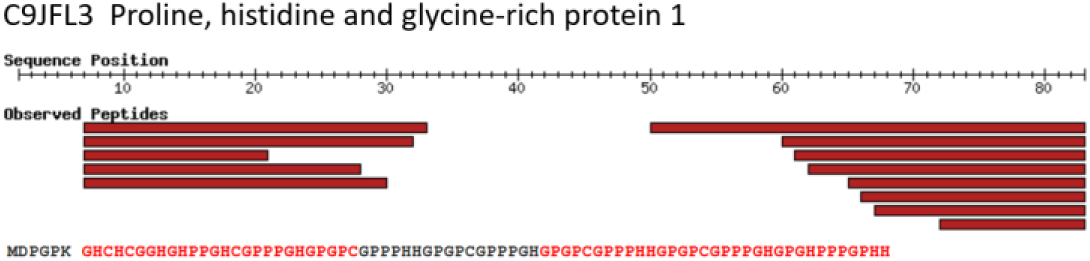
Depiction of the current status of C9JFL3, currently annotated as Proline, histidine and glycine-rich protein 1. The protein is 83 amino acids long, but produces no useful fully tryptic peptides—only one that is too short and one that is too long. Yet, due to its high abundance in some samples, many miscleaved peptides are detected, easily providing the minimum evidence. The red bars indicate well detected peptides in PeptideAtlas. Multiple semi-tryptic peptides originate from the only cleavage site after the sixth amino acid. Multiple non-tryptic peptides originate from the C terminus.

A related and difficult class of proteins is the olfactory receptors. There are four PE1 entries based on non-MS evidence, and 401 PE2,3,4 entries that are annotated as being olfactory receptors in neXtProt. None of these has the requisite 2 × 9 uniquely mapping peptides found in PeptideAtlas. Of the 401 entries, 15 do have a single peptide mapping to them. However, a manual inspection of the available spectral evidence indicates that none of these can be called a solid detection. In most cases spectra are questionable or too short to be confident about the mapping. In all, the meager evidence for olfactory receptor proteins is far more consistent with false positives than real MS detections. This is perhaps moot since the evidence as is does not meet the guidelines but serves as an important reminder that PeptideAtlas does contain some false positives, and additional stringency of multiple detections and expert review of spectra are required for high confidence. The null hypothesis therefore remains that, among all 1500+ samples collated in the PeptideAtlas build, there are zero credible detections of olfactory receptors despite some previous hints to the contrary^48^. Interestingly, if it is accepted that there have been zero credible detections of olfactory receptors via MS, one can use the putative matches to olfactory receptors in any data set to provide an independent estimate of the true FDR of the data set. In any case, after substantial discussion, the group felt that, although detection of olfactory receptors by MS has proven to be extremely refractory^49^, it should not be insurmountable, and efforts should continue. The successful detection will likely require isolation of the most appropriate olfactory cilia membrane samples, high levels of detergent to free these proteins from the membrane, enrichment with affinity reagents, and finally detection via MS of the enriched protein sample — a difficult challenge indeed. If transcript levels are extremely low and expression of any single of the ∼400 olfactory receptors is tightly limited to only one receptor in any cell at any time, detection may be not feasible due to limit of detection of current MS and possibly antibody-based methods.

The final major proposed change to the HPP pipeline is the addition of MassIVE-KB^38^ to the workflow. Whereas the current HPP pipeline (as described above and shown in Figure 1) includes only PeptideAtlas as the data set reprocessing engine, it was agreed that adding MassIVE-KB as a second reprocessing engine may have substantial benefits. While data sets reprocessed in both PeptideAtlas and MassIVE-KB have substantial overlap, this is not 100% and since MassIVE-KB has similar stringency criteria as PeptideAtlas, HPP output quality levels would be expected to be similar. Yet, it is known that there are protein detections in MassIVE-KB that meet the same criteria used by PeptideAtlas and neXtProt that should be captured by the HPP Pipeline^38^. To guard against the possible doubling of FDR by combining these resources, the HPP Pipeline will require that minimal evidence for a PE1 protein (i.e. two uniquely mapping non-nested peptides of length 9AA or more) must come from <u>either</u> PeptideAtlas <u>or</u> Massive-KB, but <u>not</u> a mixture of peptides from each. In other words, combining a singleton peptide from one resource with a singleton from the other resource will not be deemed sufficient until all evidence is reprocessed and validated by a single resource within the HPP Pipeline.

## Conclusion

As the HPP approaches one of its major initial goals (achieving credible detection of all proteins coded by the human genome), the HPP MS Data Interpretation Guidelines that served the project well since 2016 have now been clarified and enhanced with broad consensus of the HPP leadership. These revisions address some previous ambiguities that have emerged and address issues that seemed insignificant when the goal was distant. The new guidelines provide an enhanced framework for ensuring that the evidence used to substantiate future protein detection claims remains of very high quality. As such we trust they will help guide the global proteomics community on the path to missing protein discovery and functional understanding of proteins in the full biological detail of their spatiotemporal networks, pathways, molecular complexes, transport, and localization.

## Supporting information

Guidelines Checklist

## Supporting Information

Supplementary Material 1: HPP Mass Spectrometry Data Interpretation Guidelines Version 3.0 checklist and documentation.

Supplementary Table 1. Supplemental Table 1. neXtProt protein entries with only 0 or 1 uniquely mapping peptides of length 9 AA or greater using any of 5 proteases.

## Notes

The authors declare no competing financial interest.

## Acknowledgements

This work was funded in part by the National Institutes of Health grants R01GM087221 (EWD/RLM), R24GM127667 (EWD), U54EB020406 (EWD), R01HL133135 (RLM), U19AG02312 (RLM), U54ES017885 (GSO), U24CA210967-01 (GSO), R01LM013115 (NB) and P41GM103484 (NB); National Science Foundation grants ABI-1759980 (NB), DBI-1933311 (EWD), and IOS-1922871 (EWD); Canadian Institutes of Health Research 148408 (CMO); Korean Ministry of Health and Welfare HI13C2098 (YKP); French Ministry of Higher Education, Research and Innovation, ProFI project, ANR-10-INBS-08 (YV); also in part by the National Eye Institute (NEI), National Human Genome Research Institute (NHGRI), National Heart, Lung, and Blood Institute (NHLBI), National Institute of Allergy and Infectious Diseases (NIAID), National Institute of Diabetes and Digestive and Kidney Diseases (NIDDK), National Institute of General Medical Sciences (NIGMS), and National Institute of Mental Health (NIMH) of the National Institutes of Health under Award Number U24HG007822 (SO) (the content is solely the responsibility of the authors and does not necessarily represent the official views of the National Institutes of Health).

