## Supplementary material for "Human Proteome Project Mass Spectrometry Data Interpretation Guidelines 3.0": Guidelines Checklist

Version 3.0.0 – September 27, 2019

The following checklist is a brief summary of the full guidelines. This checklist must be completed by authors and submitted along with the manuscript. See pages 2-4 of this document for a more detailed description of each item in the checklist. Each item in the checklist must be checked when deemed completed, or marked as NA (Not Applicable), or NC (Not Completed); Each NA or NC *must* be explained. The second box ("Loc") must contain the location of the requested information (p12L20 for page 12 line 20, ST2 for supplementary table 2, etc.)

| **General guidelines for all manuscripts:** | | |
| --- | --- | --- |
| √ | Loc | 1. Complete this HPP MS Data Interpretation Guidelines checklist and submit with your manuscript. |
| 2. Data deposition guidelines | | |
|  |  | 2a. Deposit all MS proteomics data to a ProteomeXchange repository as a “complete” submission. |
|  |  | 2b. Include analysis reference files (search database, spectral library, transition list, etc.) in submission. |
|  |  | 2c. Provide the PXD identifier(s) in the manuscript abstract. |
|  |  | 2d. Provide the reviewer login credentials if the dataset is not yet public. |
|  |  | 3. Use the most recent version of the neXtProt reference proteome for all informatics analyses, particularly with respect to new PE1 protein detection claims. |
| 4. FDR-related guidelines | | |
|  |  | 4a. Describe in detail the calculation of FDRs at the PSM, peptide, and protein levels. |
|  |  | 4b. Report the PSM-, peptide-, and protein-level FDR values along with the total number of expected false positives at each level, using precision appropriate to the uncertainty in computed FDR. |
|  |  | 4c. Present large-scale results thresholded at equal to or lower than 1% protein-level global FDR. |
|  |  | 4d. If any large-scale datasets are individually thresholded and then combined, calculate the new, higher peptide- and protein-level FDRs for the combined result. |
| **Guidelines for claims of new PE1 protein detections (i.e., presenting evidence to categorize a protein to PE1)** | | |
|  |  | 5a. If using DDA mass spectrometry for such claims, present high mass-accuracy, high signal-to-noise ratio (SNR), and clearly annotated spectra. Scrutinize spectra for missing and extra peaks. |
|  |  | 5b. Present high mass-accuracy, high-SNR, clearly annotated spectra of synthetic peptides that match the spectra supporting the claims. Peptides from recombinant proteins are acceptable synthetics. |
|  |  | 5c. Provide Universal Spectrum Identifiers (USIs) for all natural and synthetic peptide spectra that support such claims, ideally as a supplementary data table. |
|  |  | 6. If using SRM verification for such claims, present target traces alongside synthetic heavy-labeled peptide traces, demonstrating co-elution and closely matching fragment mass intensity patterns. |
|  |  | 7. If using DIA MS, then, if the data are analyzed with XICs, apply the above SRM guidelines (6); if the data are analyzed by extracting deconvoluted spectra, apply the above DDA guidelines 5a-5c. |
|  |  | 8. Even when very high confidence peptide identifications are demonstrated, consider alternate mappings of the peptide to proteins other than the claimed one. Consider isobaric sequence/mass modification variants, all known SAAVs, and unreported SAAVs. |
|  |  | 9. Support such claims by two or more distinct uniquely-mapping, non-nested peptide sequences of length ≥9 amino acids with the above evidence in the same paper. When 2 peptides overlap, the total extent must be ≥18 amino acids. When weaker evidence is offered for such a claim, justify that other peptides cannot be expected by any common digestion proteases. When 2 or more proteins are *exactly* sequence identical (irrespective of SAAVs), peptides are considered uniquely mapping if they map only to the group, and such proteins will have the same PE level and be counted. |

Author comments (use this space and extra pages to explain any nonadherence [NA/NC] in the above checklist):

(see extended description for each of the above items on pages 2 - 4 below)

**Extended Detail on Checklist items:**

The following pages provide some additional detail on the intentionally terse one-page checklist. Users new to this version of the HPP MS Data Interpretation Guidelines should read these extended descriptions before using the checklist.

1. **Complete this HPP MS Data Interpretation Guidelines checklist and submit with your manuscript**. Page 1 of this document must be submitted as supplementary material for the editor, reviewers, and readers. The completed checklist is required before a manuscript will be sent to reviewers. Each item in the checklist must be either checked, marked as NA (Not Applicable), or marked as NC (Not Completed). Please explain NA or NC entries or any other variances in the Author Comments section at the bottom of the checklist. In the second column, enter the location where the requested information may be found (e.g., p12 for page 12, p12L10 for page 12 line 10, ST2 for supplementary table 2, etc.) This will assist reviewers and readers in finding the information quickly. Manuscripts received without a checklist will be returned without review.
2. **Guidelines for data repository deposition**
   a. **Deposit all MS proteomics data to a ProteomeXchange repository as a complete submission.** All depositions are required to be “Complete” submissions instead of “Partial” submissions. ProteomeXchange deposition must be completed prior to submission of the manuscript to the journal. Synthetic peptide MS runs must also be deposited and clearly marked as such.
   b. **Include analysis reference files (search database, spectral library, transition list, etc.) in submission.** Include all supplemental data files used in the analysis. Included software parameter files if relevant.
   c. **Provide the PXD identifier(s) in the manuscript abstract.**
   d. **Provide the reviewer login credentials if the dataset is not yet public**. Reviewer login information at the repository must be provided in the manuscript if the dataset is not already publicly released.
3. **Use the most recent version of the neXtProt reference proteome for all informatics analyses, particularly with respect to new PE1 protein detection claims.** Informatics analysis should always be presented in comparison with the most recent proteome references, rather than older versions thereof. For the HPP special issues, the required version will be listed in the call for papers, usually a January or February release.
4. **FDR-related guidelines**
   a. **Describe in detail the calculation of FDRs at the PSM, peptide, and protein levels**. Describe which tools are used to estimate the false discovery rate (FDR) at the peptide-spectrum-match (PSM) level, at the distinct peptide sequence level, and at the protein level. Briefly describe the approach and what assumptions are made or implied, and any corrections for the fraction of the proteome covered. If you use novel or uncommon tools and criteria, compare your results with results with tools that are widely used in the community.
   b. **Report the PSM-, peptide-, and protein-level FDR values along with the total number of expected false positives at each level, using precision appropriate to the uncertainty in computed FDR**. Report the actual numbers of true positives and false positives at each level based on the thresholds used. Do not report the FDR with many significant digits since all current FDR calculation methods have substantial uncertainties.
   c. **Present large-scale results thresholded at equal to or lower than 1% protein-level global FDR**. The 1% is somewhat arbitrary but well accepted and remains set as the upper limit. For many datasets from modern instrumentation, achieving a 1% global FDR may include very low-quality results with a **local** FDR worse than 10%, which is undesirable. A global FDR lower than 1% is encouraged, but it should never be higher than 1%. Similarly, PSMs and peptides with a local FDR worse than 10% should not be included. The common mistake of thresholding at 1% FDR and then assuming that all surviving results are correct, no matter how surprising, must be avoided.
   d. **If any large-scale datasets are individually thresholded and then combined, calculate the new, higher peptide- and protein-level FDRs for the combined result**. When datasets are combined, the true positives will mostly overlap, while the false positives will be scattered randomly across the proteome and thus overlap far less. This means that the FDR will be higher in the combined dataset.

Whereas the above guidelines apply to all manuscripts presenting mass spectrometry data, the following guidelines apply only to manuscripts that are presenting evidence to promote proteins that are not currently listed in neXtProt as PE1 protein to PE1 status. This may apply to one of the “missing proteins”, which are currently in neXtProt with PE2-4. This may apply to a currently PE5 protein, although most of these entries are thought to be pseudogenes and extra care must be applied to justify that the detections are not merely variation of the common PE1 protein that the PE5 protein closely resembles. Finally, this may apply to a protein not yet listed in neXtProt, such as a lncRNA or a smORF or some other novel coding element. Care should be taken to see if the protein already exists in UniProtKB/TrEMBL or RefSeq and needs to be manually transferred to UniProtKB/Swiss-Prot and thus neXtProt. If this is the case, it is recommended that you first contact the UniProt team with your evidence to request curation of this protein into UniProtKB/Swiss-Prot. The UniProt curation team uses several different evidence sources, including the HPP MS Data Interpretation Guidelines to manually assign the PE level when an entry is moved into UniProtKB/Swiss-Prot, whereas the PE level for proteins in UniProtKB/TrEMBL is assigned automatically, again using criteria including the HPP guidelines.

5. **Guidelines for data-dependent acquisition (DDA) MS datasets**
   a. **If using DDA mass spectrometry for such claims, present high mass-accuracy, high signal-to-noise ratio (SNR), and clearly annotated spectra. Scrutinize spectra for missing and extra peaks**. Annotated spectra (i.e., spectra with the matched peaks clearly labeled) must be provided in the supplementary material for the manuscript. While low mass-accuracy and low SNR spectra can still be useful for many experiments, they are not acceptable for claims of new PE1 protein detections. Time-of-flight, FT-ICR, and Orbitrap-type instruments are considered in these guidelines as having high mass accuracy (when properly calibrated) in these guidelines. The spectra should be examined closely to determine if there are peaks missing that should be expected, if there are peaks present that are unexplained, and if a small alteration of the putative sequence would yield a much better match. This may indicate a false positive of a kind that is not modeled well by decoys.
   b. **Present high mass accuracy, high-SNR, clearly annotated spectra of synthetic peptides that match the spectra supporting the claims. Peptides from recombinant proteins are acceptable synthetics.** Synthetic peptides are powerful tools for determining the correct identification of spectra. For each PSM corresponding to claim of a new PE1 protein, compare that PSM with a synthetic peptide (or recombinant protein product) spectrum of the same ion. All the major ions should have closely matching intensities in both spectra. If generating new reference spectra, it is encouraged to use the same high mass-accuracy instrument to verify matching intensity patterns and retention times. Closely matching spectra of the same peptidoform ion (same modifications and charge) from SRMAtlas, ProteomeTools, or similar resources is acceptable. Predicted spectra may be used as part of analysis workflows, but not as comparison reference spectra.
   c. **Provide Universal Spectrum Identifiers (USIs) for all natural and synthetic peptide spectra that support such claims, ideally as a supplementary data table.** The USI provides a mechanism to uniquely identify a spectrum being held up as evidence for an important claim. The USI will allow readers to access these important spectra in public data repositories in order to discuss correctness of the claims. See <http://psidev.info/USI> for more information.
6. **If using SRM verification for such claims, present target traces alongside synthetic heavy-labeled peptide traces, demonstrating co-elution and closely matching fragment mass intensity patterns**. All SRM runs performed must have spiked-in heavy labeled peptides corresponding to the putative identifications. The heavy-labeled peptides should be spiked in at an abundance similar to the target peptides so that minor impurities in the reference do not contribute to the target signal. Annotated chromatograms must be provided in the supplementary material of the manuscript. Solid peptide sequence evidence does not alter the uncertainties in matching that peptide uniquely to a protein (guideline 8). This guideline may also apply to PRM traces, although since PRM generates full MS/MS spectra, Guideline 5a-5c may be applied to PRM data instead. Guidelines 8 and 9 also apply for SRMs.
7. **If using DIA MS, then, if the data are analyzed with XICs, apply the above SRM guidelines (6); if the data are analyzed by extracting deconvoluted spectra, apply the above DDA guidelines 5a-5c.** DIA-MS workflows such as SWATH-MS or the equivalent on other instrument types yield highly multiplexed spectra that make confident identification of peptides difficult. The guidelines that apply depend on the data analysis strategy. If the data are analyzed via extracted ion chromatograms (XICs) such as with OpenSWATH, Spectronaut, PeakView, etc. then the SRM guideline 6 above applies. If the data are analyzed via extracted deconvoluted spectra such as with DIA-Umpire or DISCO, then the DDA Guideline 5a-5c above applies. In addition to the raw data, the extracted deconvoluted spectra must also be submitted to ProteomeXchange repository.
8. **Even when very high confidence peptide identifications are demonstrated, consider alternate mappings of the peptide to proteins other than the claimed one. Consider isobaric sequence/mass modification variants, all known SAAVs, and unreported SAAVs**. Even when a peptide identification is shown to be very highly confident, care should be taken when mapping it to a protein or novel coding element. Consider whether I=L, N[Deamidated]=D, Q[Deamidated]=E, GG=N, Q≈K, F≈M[Oxidation], or other isobaric or near isobaric substitutions could change the mapping of the peptide from an extraordinary result to a mapping to a commonly-observed protein. Consider if a known single amino-acid variation (SAAV) in neXtProt could turn an extraordinary result into an ordinary result. Consider if a single amino-acid change, not yet annotated in a well-known source, could turn an extraordinary result into a questionable result. Check more than one reference proteome (e.g., RefSeq may have entries that UniProt and Ensembl do not, and vice versa). A tool to assist with this analysis is available at neXtProt at <https://www.nextprot.org/tools/peptide-uniqueness-checker> (Unicity Checker), and another at PeptideAtlas at <http://peptideatlas.org/map> (ProteoMapper).
9. **Support such claims by two or more distinct uniquely-mapping, non-nested peptide sequences of length ≥9 amino acids** **with the above evidence in the same paper. When 2 peptides overlap, the total extent must be ≥18 amino acids. When weaker evidence is offered for such a claim, justify that other peptides cannot be expected by any common digestion proteases. When 2 or more proteins are *exactly* sequence identical (irrespective of SAAVs), peptides are considered uniquely mapping if they map only to the group, and such proteins will have the same PE level and be counted**. Single-peptide detections simply have too high a chance of being some type of pernicious false positive to be sufficient for claiming a new PE1 protein detection. Likewise, short peptides of length 8 or smaller have relatively few peaks and have an increased chance of mapping to immunoglobulins or other sequences not readily apparent in the reference proteome. Nested peptides (where one sequence is fully subsumed within another) do provide additional confidence that the peptide identification is correct, but provide no additional evidence that the peptide-to-protein mapping is unique. In rare cases only a single uniquely mapping peptide can be generated even when applying different proteases; this may then be sufficient if the case is well justified. If the entire mature form of a very short protein has 100% coverage with excellent spectra but yet does not strictly meet the guidelines, this may indicate a clear example of a justifiable exception. The practice of offering a single new suitable peptide to complement a pre-existing different suitable peptide already in PeptideAtlas and neXtProt is permitted, but the PeptideAtlas peptide spectrum must also be scrutinized and compared with a synthetic peptide spectrum in accordance with the above guidelines with all evidence presented in the paper. Alternatively, if it is desirable to present evidence that does not meet these criteria for new PE1 protein detection claims, the implicated proteins may be offered as “candidate detections” to enable capture of this information by other researchers for follow up by further experiments. A special HPP PE classification exceptions review panel is being established with membership from HUPO’s HPP, PeptideAtlas, and neXtProt to evaluate “exceptional” PE categorization cases.
